## supplemental Figure 1-12 for "Prdm1 Positively Regulates Liver Group 1 ILCs Cancer Immune Surveillance and Preserves Functional Heterogeneity"

### Affiliations

### Conflict-of-interest statement

The authors have declared that no conflict of interest exists.

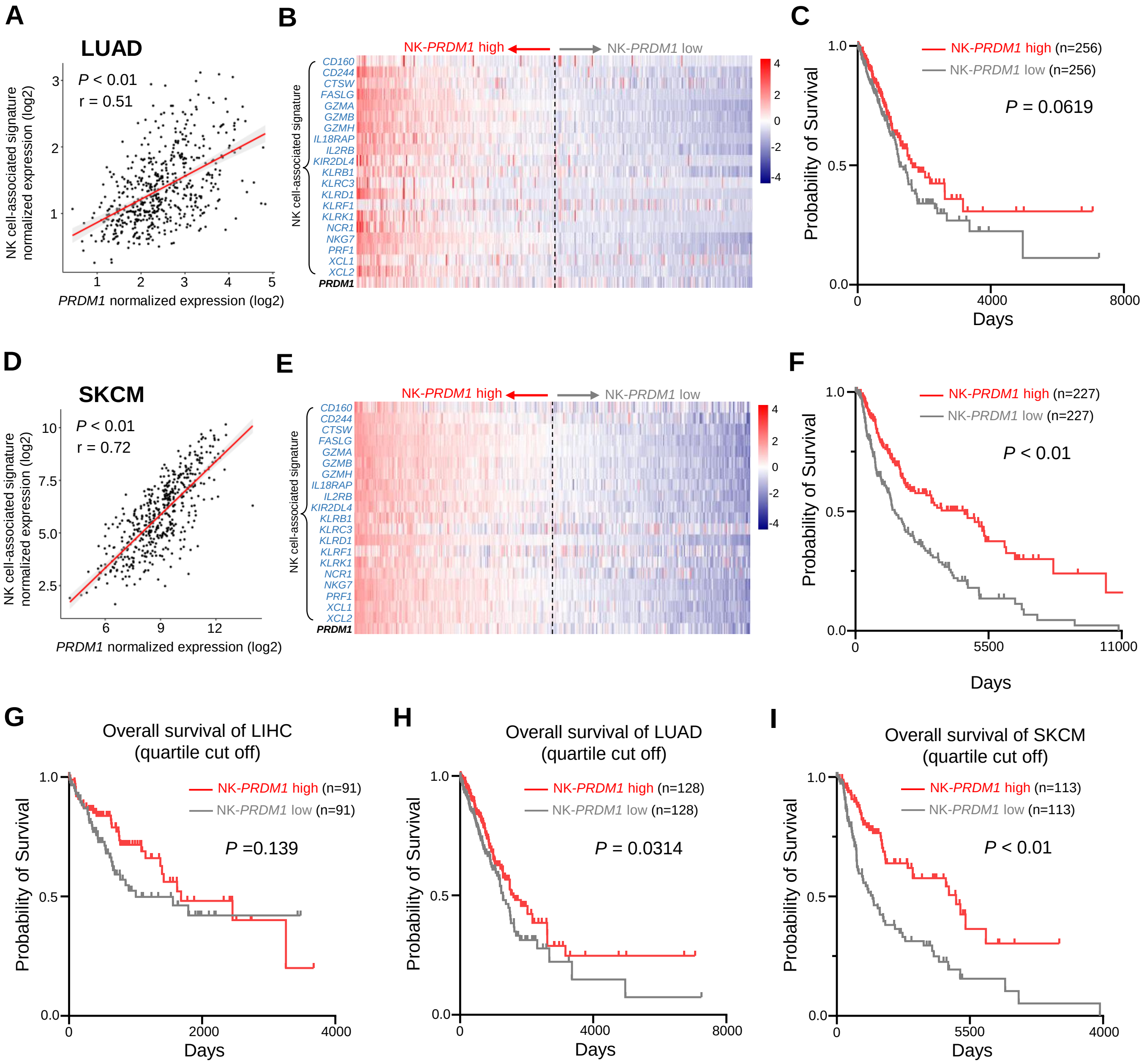


**Supplemental Figure 1. High expression of *PRDM1-NK* signature predicts better overall survival of cancer patients.**
(**A and D**) Correlation between the average expression of NK cell-associated signature and *PRDM1* in lung adenocarcinoma (**A**) (LUAD, n=497) and skin cutaneous melanoma (SKCM, n=454) (**D**) patients from TCGA datasets. (**B and E**) Heatmap of the ordered, z-score normalized expression values for *PRDM1* and NK cell-associated genes in LUAD (**B**) and SKCM (**E**) patients. High and low expression of *NK-PRDM1* signature are indicated. (**C and F**) Prognostic value of the *NK-PRDM1* signature for overall survival of LUAD (**C**) and SKCM (**F**) patients comparing high and low samples with a median cutoff. (**G-I**) Prognostic value of the *NK-PRDM1* signature for overall survival of LIHC (**G**), LUAD (**H**), and SKCM (**I**) patients comparing high and low samples with a quartile cutoff. *P*, *P*-value; r, pearson correlation coefficient.

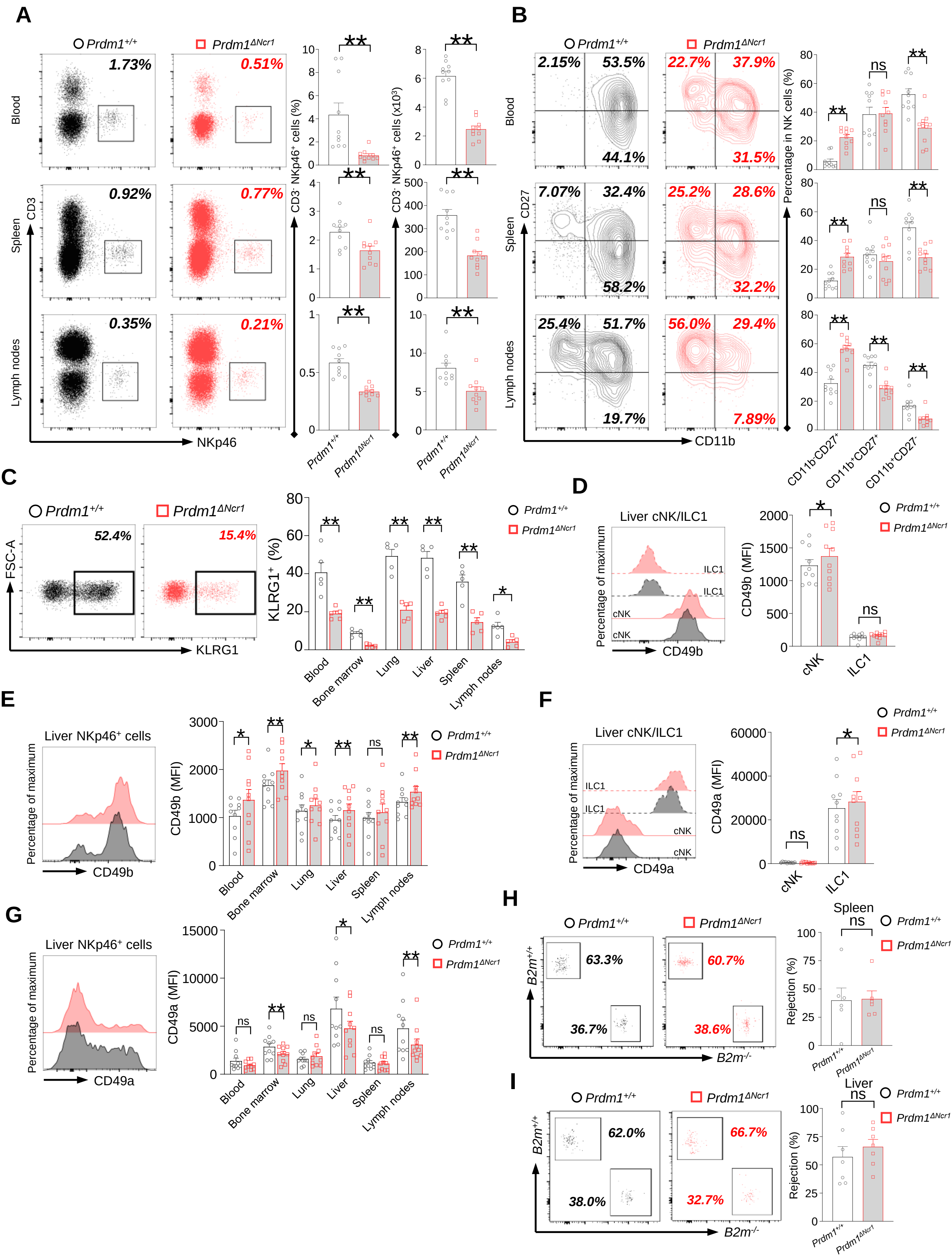


**Supplemental Figure 2. *Prdm1* plays an essential role in group 1 ILCs homeostasis and maturation.**
(**A**) Representative flow cytometric plots (left) and quantification (right) of the proportion and absolute number of $CD45^+CD3^-NKp46^+$ cells among lymphocytes in liver, lung, and bone marrow (n=10). (**B**) Representative flow cytometric plots (left) of the CD11b and CD27 expression within $CD3^-NK1.1^+NKp46^+CD49b^+$ cells in liver, lung, and bone marrow (n=10). Right panel showed the percentage of distinct maturation stages of NK cells. **(C)** Representative flow cytometry analyses of $KLRG1^+$ cells in $CD3^-NK1.1^+NKp46^+CD49b^+$ cells in liver, and their quantification in blood, bone marrow, lung, liver, spleen, and lymph nodes (n=5). (**D-G**) Representative flow cytometry plots of the mean fluorescence intensities (MFIs) of CD49b and CD49a in liver cNK cells and ILC1s (**D** and **F**), and in $NKp46^+$ cells in blood, bone marrow, lung, liver, spleen, and lymph nodes (**E** and **G**) (n=10). (**H** and **I**) Splenocytes from *B2m*$^{-/-}$ and *B2m*$^{+/+}$ were labeled with CFDASE and eF670 respectively. Labeled cells were 1:1 mixed and injected i.v. into *Prdm1*$^{+/+}$ and *Prdm1*$^{\Delta Ncr1}$ recipient mice to evaluate *in vivo* NK cell target-killing ability. Representative flow cytometric plots (left) of transferred cells recovered from recipient mice and percentage (right) of NK cell-specific rejection of donor cells in spleen (n=6) (**H**) and liver (n=7) (**I**) between *Prdm1*$^{+/+}$ and *Prdm1*$^{\Delta Ncr1}$ mice. Data are presented as the mean±SEM and were analyzed by 2-tailed, paired t-test. Differences were evaluated between littermates. Each circle and square on graphs represents an individual mouse; *P*, *P*-value; *, *P*<0.05; **, *P*<0.01, ns, not significant.

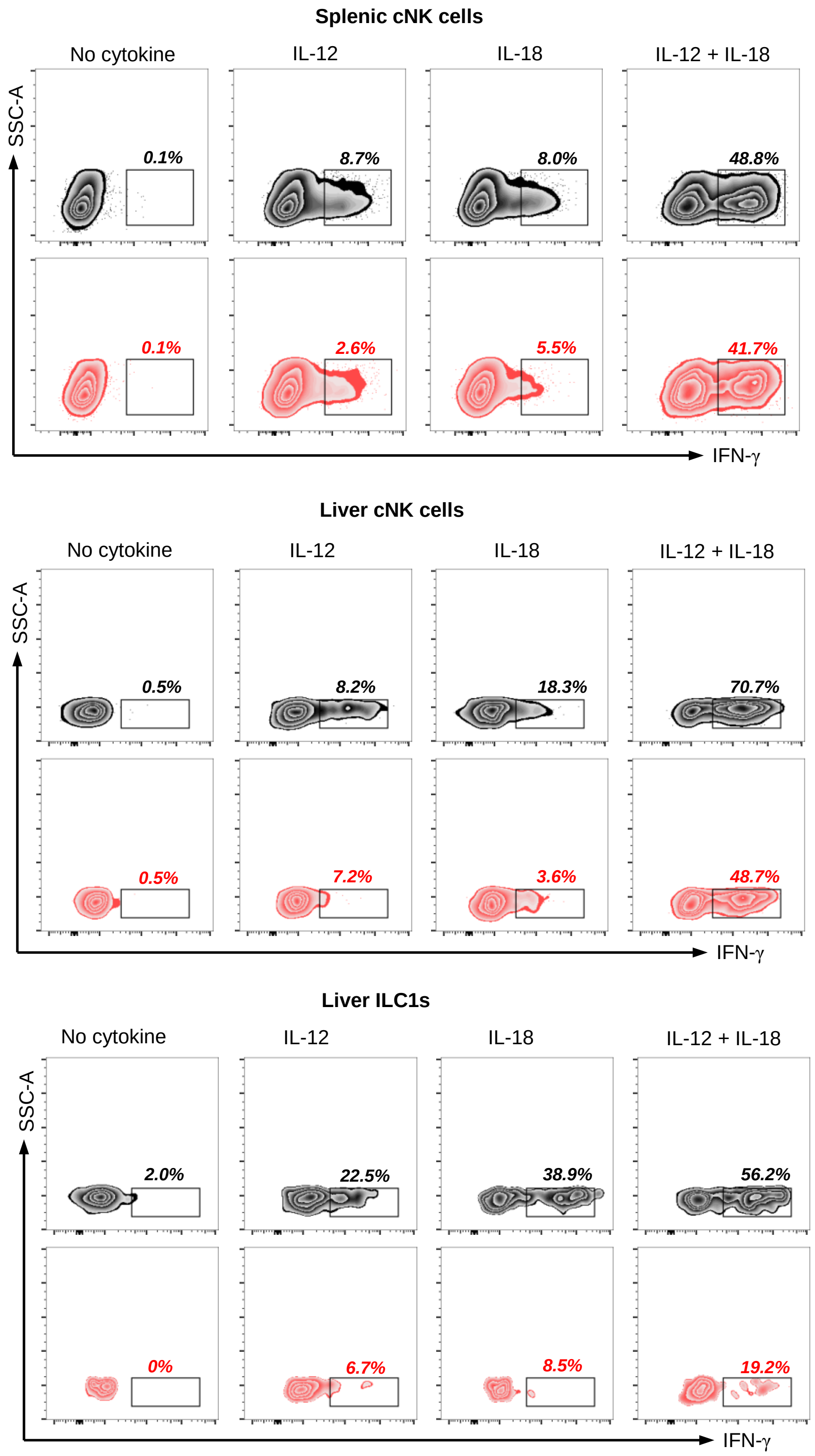


**Supplemental Figure 3. *Prdm1* affects the IFN-γ secretion ability of type I ILCs.**
Representative flow cytometric plot showing the frequency of IFN-$\gamma^+$ cNK cells and ILC1s in spleen and liver between *Prdm1*$^{+/+}$ and *Prdm1*$^{\Delta Ncr1}$ mice (n=5). Liver cells and splenocytes were costimulated in the presence or absence of IL-12 and IL-18 for 12 hours. GolgiStop was added 4 hours before intracellular staining of IFN-γ. Data are presented as the mean±SEM and were analyzed by 2-tailed, paired t-test. Differences were evaluated between littermates. Each circle and square on graphs represents an individual mouse; *P*, *P*-value; *, *P*<0.05; **, *P*<0.01, ns, not significant.

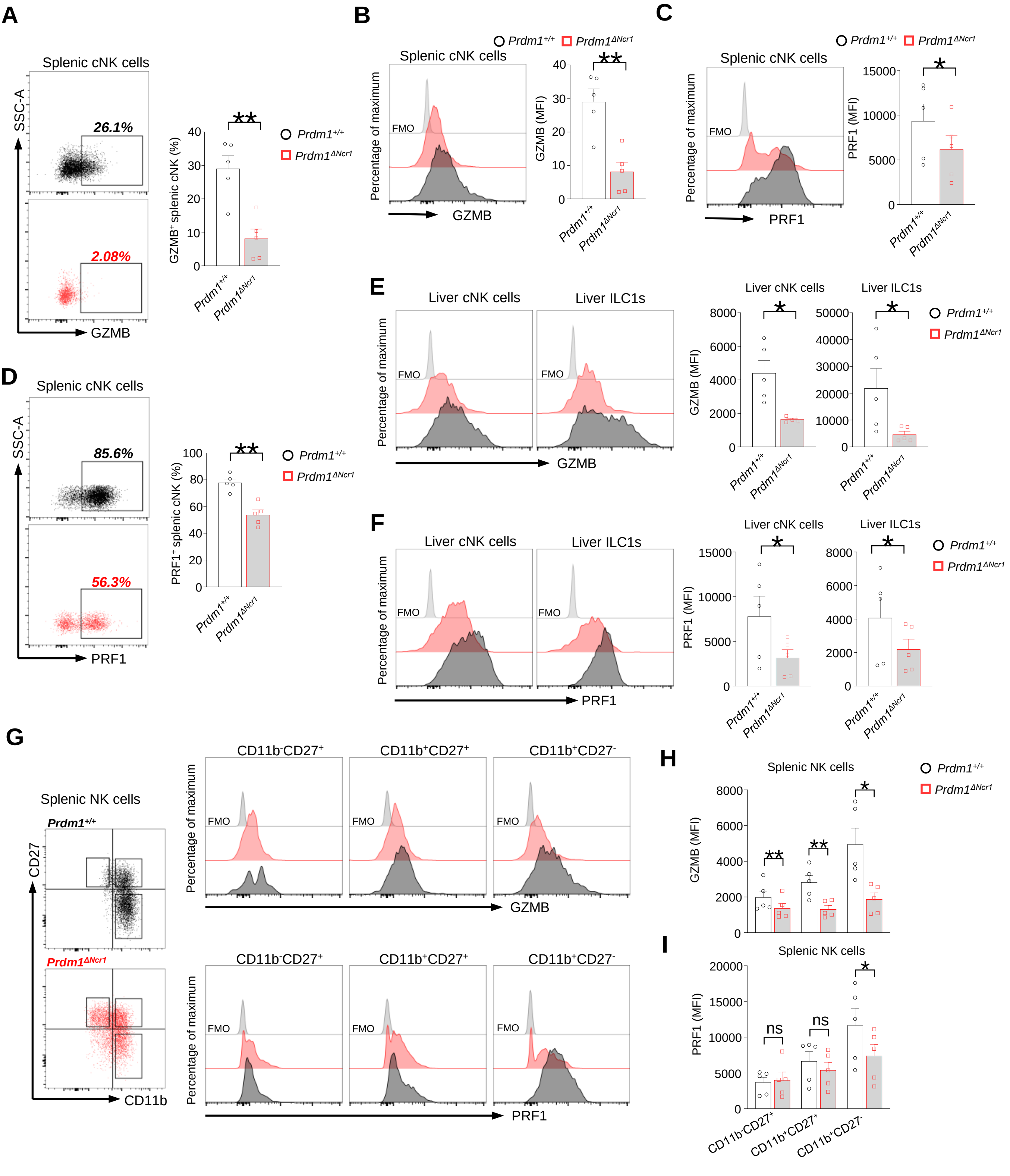

**Supplemental Figure 4. *Prdm1* deficiency impairs production of Granzyme B and Perforin in group 1 ILC.**
(**A-D**) Representative flow cytometric plot (left) and summary data (right) showing the proportion of GZMB$^+$ (**A**) and PRF1$^+$ (**D**) splenic cNK cells and relative mean fluorescence intensities (MFIs) of GZMB (**B**) and PRF1 (**C**) in splenic NK cells from *Prdm1*$^{+/+}$ and *Prdm1*$^{\Delta Ncr1}$ mice (n=5). (**E** and **F**) ) Representative flow cytometric plots (left) and cumulative data (right) showing the relative MFIs of GZMB (**E**) and PRF1 (**F**) in liver cNK cells and ILC1s from *Prdm1*$^{+/+}$ and *Prdm1*$^{\Delta Ncr1}$ mice (n=5). (**G-I**) Splenic NK cells were divided into different stages according to the expression of maturation markers CD11b and CD27. Representative flow cytometric plot (**G**) and showing the MFIs of GZMB (**H**) and PRF1 (**I**) in evaluated between littermates. Data are presented as the mean±SEM and were analyzed by 2-tailed, paired t-test. Differences were evaluated between littermates. Each circle and square on graphs represents an individual mouse; *P*, *P*-value; r, pearson correlation coefficient; *, $P$<0.05; **, $P$<0.01, ns, not significant.

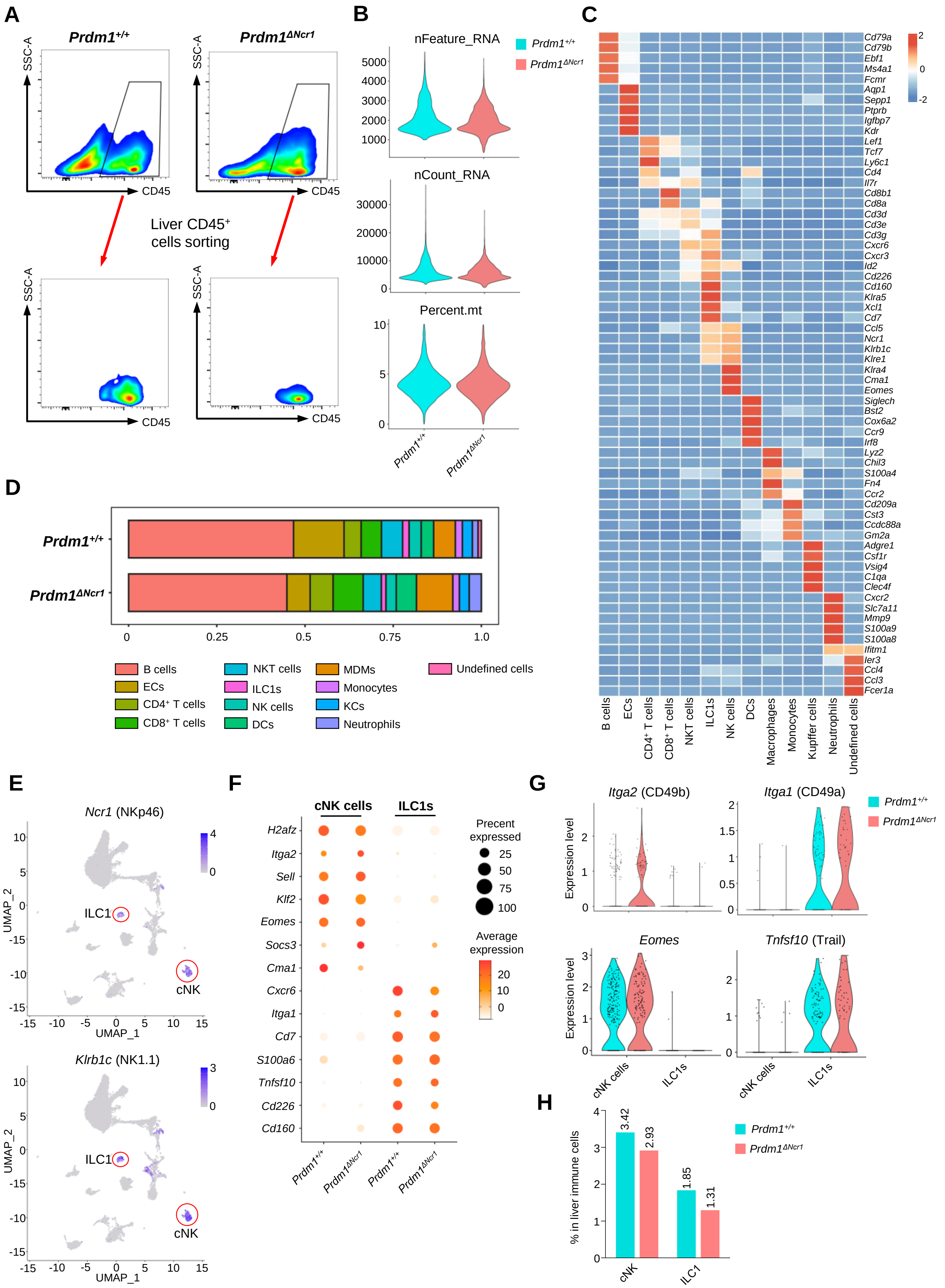


**Supplemental Figure 5. scRNA-seq identified subsets of liver CD45⁺ cell from *Prdm1*$^{+/+}$ and *Prdm1*$^{\Delta Ncr1}$ mice.**
(**A**) Liver CD45⁺ cells from *Prdm1*$^{+/+}$ and *Prdm1*$^{\Delta Ncr1}$ mice were FACS-sorted for scRNA-seq. (**B**) Quality control of the scRNA-seq data. Violin plot showing the nFeature_RNA, nCount_RNA, and percentage of mitochondrial genes of sequencing cells. nFeature_RNA represents the number of genes detected in each cell, and nCount_RNA represents the total number of molecules detected within a cell. (**C**) Heatmap showing the expression of top 5 upregulated DEGs of each cell cluster by log fold change (computed using Wilcox test in the "FindAllMarkers" function of Seurat, avg_log2FC>0.25; P<0.05). (**D**) Population of twelve liver CD45⁺ cell clusters between *Prdm1*$^{+/+}$ and *Prdm1*$^{\Delta Ncr1}$ mice. (**E**) Feature plots showing the normalized expression of *Ncr1* (NKp46) and *Klrbc1* (NK1.1) for different cell clusters. (**F**) Dot plot showing the normalized expression of marker genes in liver cNK cells and ILC1s clusters between *Prdm1*$^{+/+}$ and *Prdm1*$^{\Delta Ncr1}$. The dot size represents the percentage of cells expressing selected genes, and color intensity represents the average expression. (**G**) Violin plots showing the expression of marker genes *Itga2* (CD49b), *Itga1* (CD49a), *Eomes, and Tnfsf10* (Trail) between liver cNK cells and ILC1s between *Prdm1*$^{+/+}$ and *Prdm1*$^{\Delta Ncr1}$ mice. (**H**) The percentage of liver cNK cells and ILC1s in liver immune cells between *Prdm1*$^{+/+}$ and *Prdm1*$^{\Delta Ncr1}$.

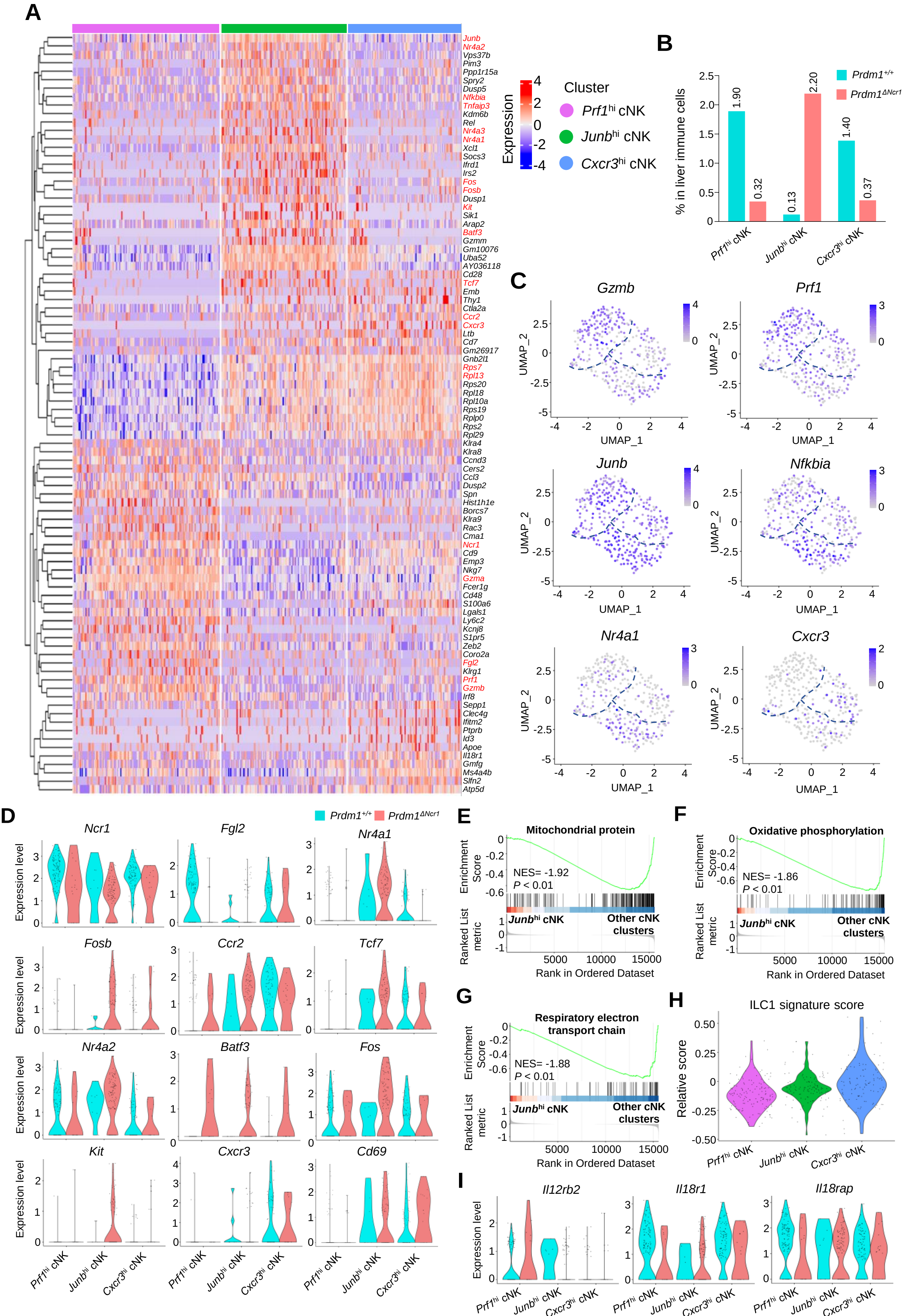


**Supplemental Figure 6. Cluster-specific markers of liver cNK cell clusters.**
(**A**) Heatmap showing the expression of top 30 upregulated DEGs by log fold change (computed using Wilcox test in the "FindAllMarkers" function of Seurat, avg_log2FC>0.25; P<0.05), across the three liver cNK cell clusters (*Prf1$^{hi}$* cNK, *Junb$^{hi}$* cNK, *Cxcr3$^{hi}$* cNK) within *Prdm1$^{+/+}$* and *Prdm1$^{\Delta Ncr1}$* mice. (**B**) The percentage of *Prf1$^{hi}$* cNK, *Junb$^{hi}$* cNK, and *Cxcr3$^{hi}$* cNK cells in liver immune cells between *Prdm1$^{+/+}$* and *Prdm1$^{\Delta Ncr1}$*. (**C**) Feature plots showing the normalized expression of selected markers for liver cNK cell populations. (**D**) Violin plots showing the normalized expression of DEGs for each cNK cluster within *Prdm1$^{+/+}$* and *Prdm1$^{\Delta Ncr1}$* mice. (**E-G**) GSEA of the enrichment of Mitochondrial protein (**E**), Oxidative phosphorylation (**F**), and Respiratory electron transport chain (**G**) in *Junb*$^{hi}$ cNK cluster compared with clusters of *Prf1*$^{hi}$ and *Cxcr3*$^{hi}$ cNK cells. NES, normalized enrichment score. (**H**) Violin plot showing the ILC1 signature score for different cNK cell clusters, calculated using the signature genes of ILC1 cluster. (**I**) Violin plots showing the normalized expression of cytokine receptor genes (*Il12rb2*, *Il18r1*, and *Il18rap*) for each cNK cluster within *Prdm1$^{+/+}$* and *Prdm1$^{\Delta Ncr1}$* mice.

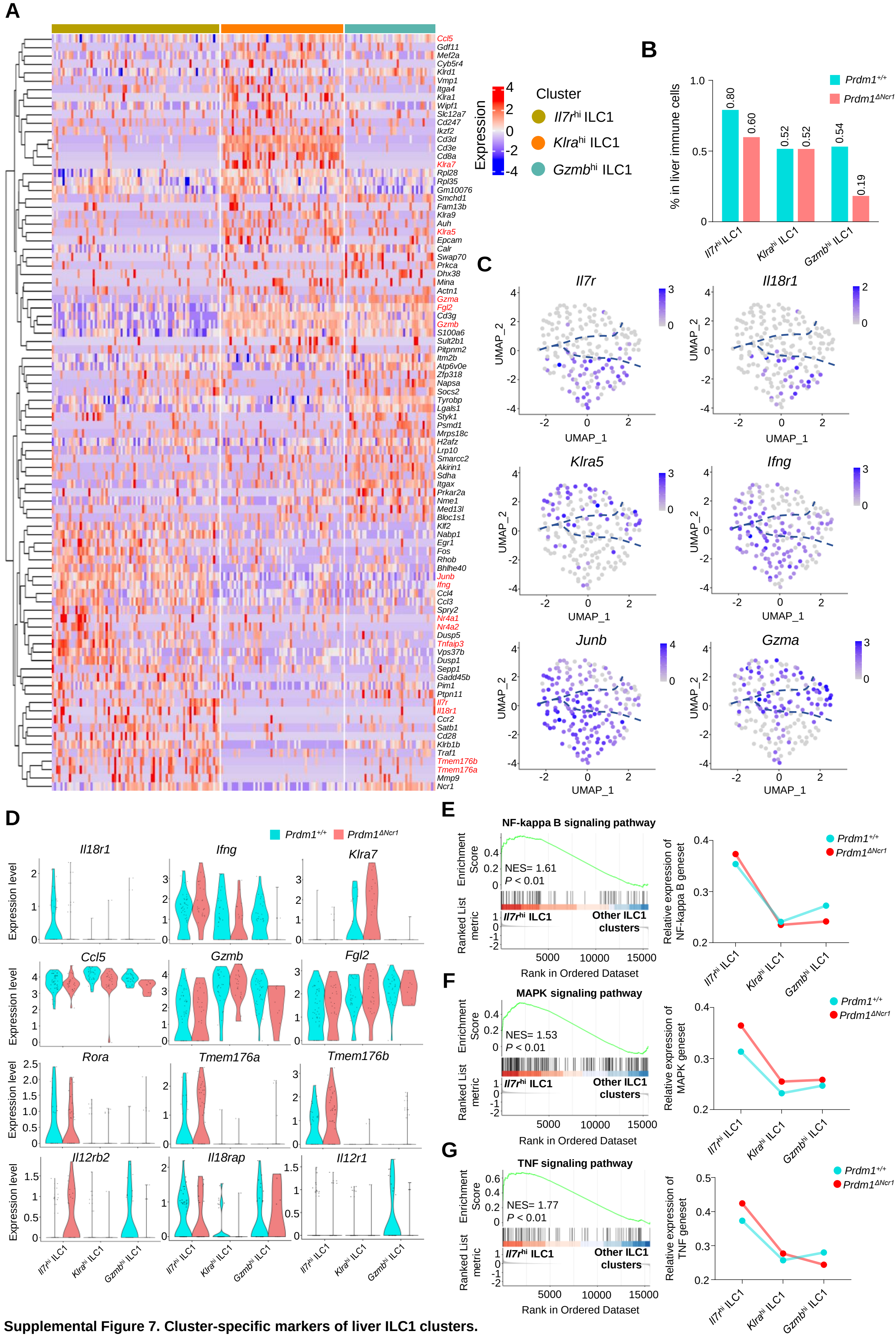


**Supplemental Figure 7. Cluster-specific markers of liver ILC1 clusters.**
(**A**) Heatmap showing the expression of top 30 upregulated DEGs by log fold change (computed using Wilcox test in the "FindAllMarkers" function of Seurat, avg_log2FC>0.25; P<0.05), across the three ILC1 clusters (*Il7r*$^{hi}$ ILC1, *Klra*$^{hi}$ ILC1, *Gzma*$^{hi}$ ILC1) within *Prdm1*$^{+/+}$ and *Prdm1*$^{\Delta Ncr1}$ mice. (**B**) The percentage of *Il7r*$^{hi}$ ILC1, *Klra*$^{hi}$ ILC1, and *Gzma*$^{hi}$ ILC1 in liver immune cells between *Prdm1*$^{+/+}$ and *Prdm1*$^{\Delta Ncr1}$. (**C**) Feature plots showing the normalized expression of selected markers for liver ILC1 populations. (**D**) Normalized expression of DEGs for each ILC1 cluster within *Prdm1*$^{+/+}$ and *Prdm1*$^{\Delta Ncr1}$ mice. (**E-G**) GSEA plots (left) depicting the enrichment of NF-kappa B (**E**) , MAPK (**F**), and TNF (**G**) signaling pathways) in *Il7r*$^{hi}$ ILC1 cluster compared with clusters of *Klra*$^{hi}$ and *Gzma*$^{hi}$ ILC1s. Right panel showed dynamic relative expression of the given gene sets from cluster1 to cluster3 between *Prdm1*$^{+/+}$ and *Prdm1*$^{\Delta Ncr1}$. Dots represent the average expression of given gene set in each cell, which was calculated through the sum of normalized expression of each individual gene within the designated gene set in every single cell. NES, normalized enrichment score.

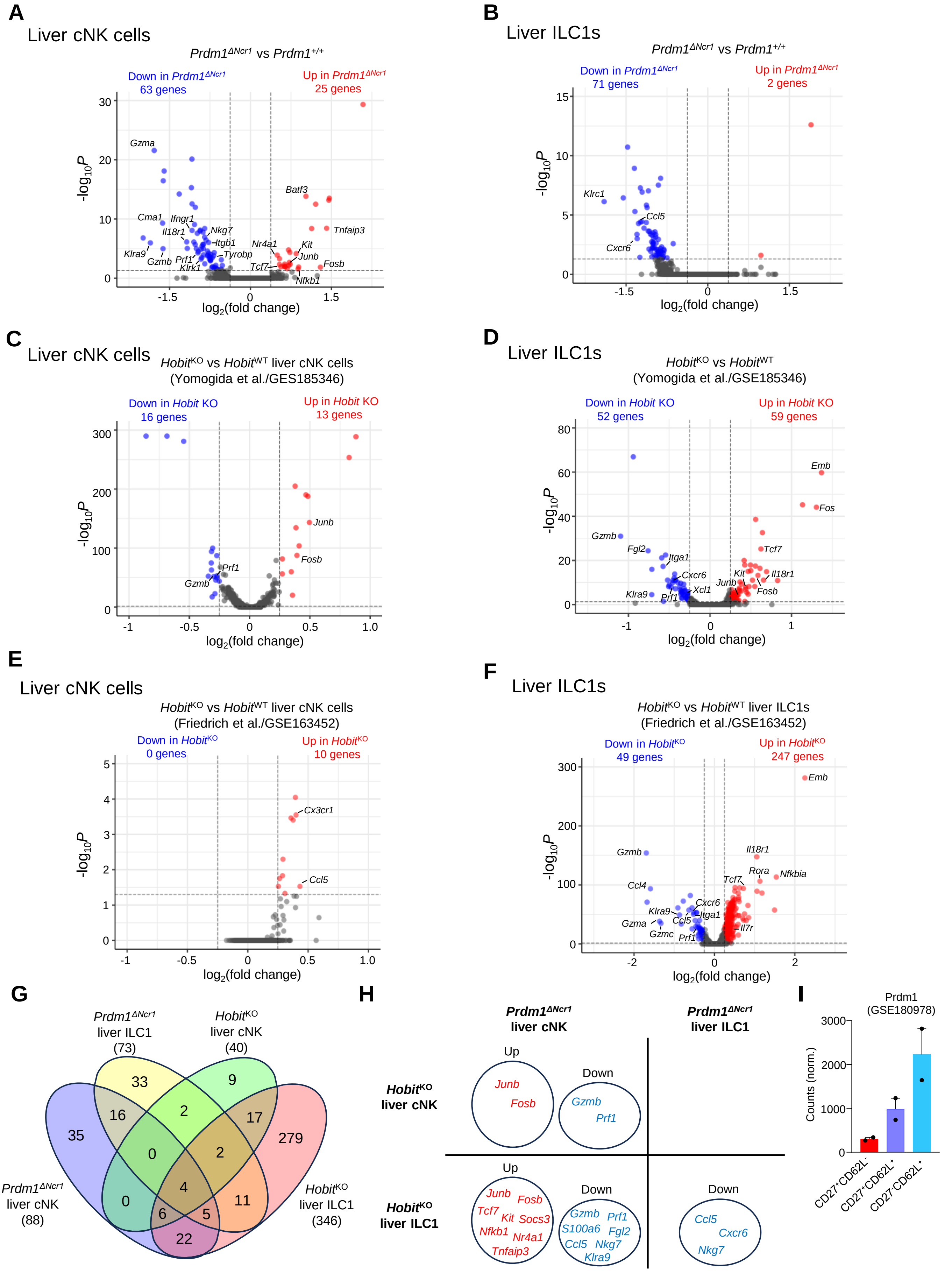

**Supplemental Figure 8. Similar and unique gene expression patterns in group 1 ILCs regulated by *Prdm1* and *Hobit*.**
**(A and B)** Volcano plots showing the DEGs (avg_log2FC>0.25; P<0.05) in *Prdm1*$^{\Delta Ncr1}$ liver cNK cells (**A**) and ILC1s (**B**) compared to *Prdm1*$^{+/+}$ samples. (**C-F**) *Hobit*$^{KO}$ liver cNK cell and ILC1 clusters were identified from published scRNA-seq data (GES185346 and GSE163452). Volcano plots showing the DEGs (avg_log2FC>0.25; P<0.05) in *Hobit*$^{KO}$ liver cNK cells (**C** and **E**) and ILC1s (**D** and **F**) in comparisons within *Hobit*$^{WT}$ clusters. (**G**) Venn diagram showing the overlapped DEGs within liver cNK cells and in *Prdm1*$^{\Delta Ncr1}$ and *Hobit*$^{KO}$ compared to control samples. (**H**) Selected DEGs common regulated in *Prdm1*$^{\Delta Ncr1}$ and *Hobit*$^{KO}$ liver cNK cells or ILC1s. Upregulated genes (red) and downregulated genes (blue) were indicated. (**I**) Bar plot showing mRNA normalized counts of *Prdm1* in specific NK cell population based on the published bulk RNA-seq data (Flommersfeld et al., GSE180978).

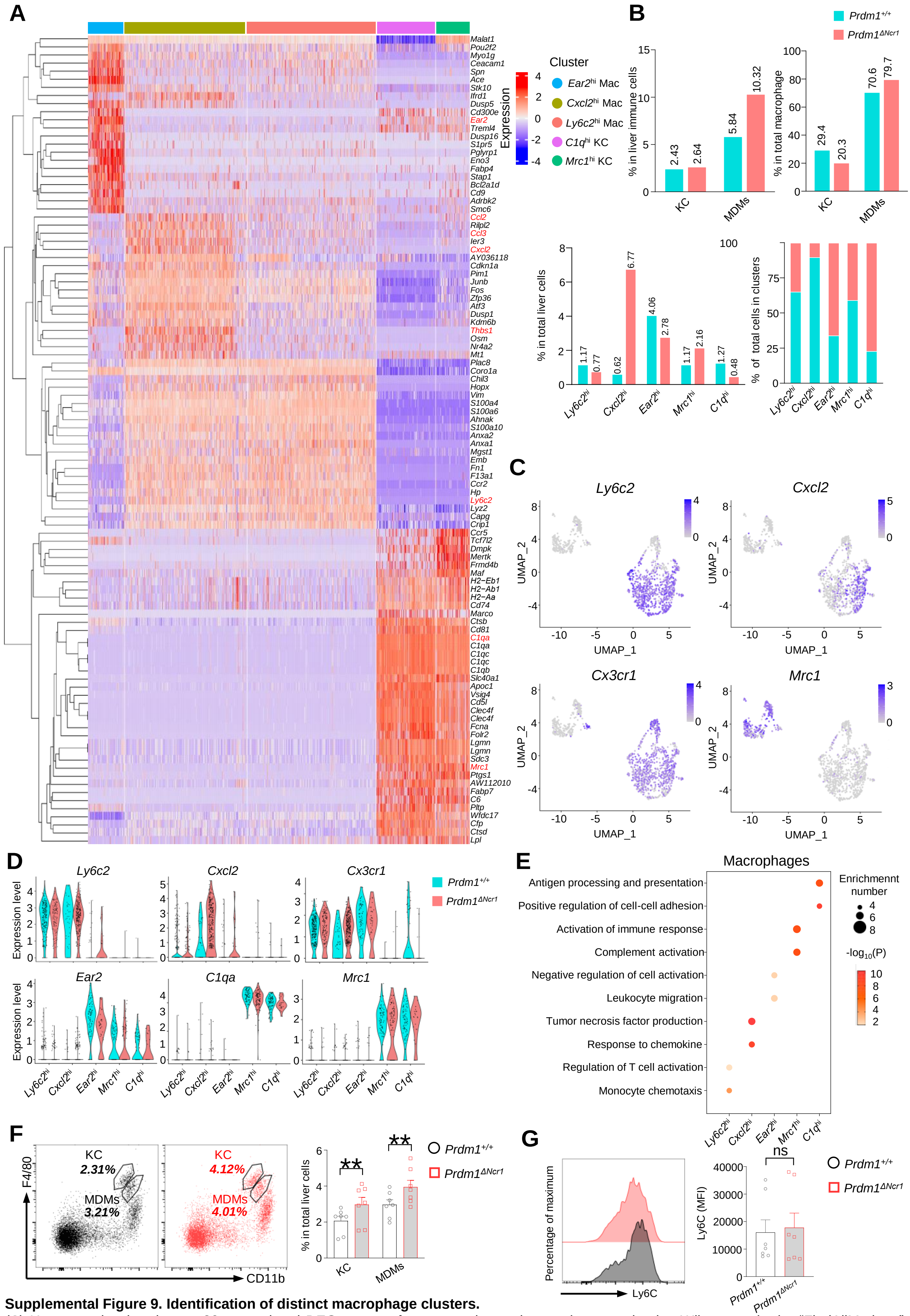


**Supplemental Figure 9. Identification of distinct macrophage clusters.**

(**A**) Heatmap showing the top 20 upregulated DEGs across four macrophage clusters (computed using Wilcox test in the "FindAllMarkers" function of Seurat, avg_log2FC>0.25; P<0.05). (**B**) Proportions of macrophages (including monocyte-derived macrophages (MDMs) and Kupffer cells (KCs) cluster) in liver immune cells (top, left). Proportions of KCs and MDMs in total macrophages (top, right).The percentage of distinct macrophage clusters in liver total macrophages (bottom, left) and within clusters (bottom, right) between *Prdm1*$^{+/+}$ and *Prdm1*$^{\Delta Ncr1}$. (**C**) Feature plots showing the normalized expression of selected markers for macrophage populations. (**D**) Violin plots showing the expression of selected markers of each macrophage cluster within *Prdm1*$^{+/+}$ and *Prdm1*$^{\Delta Ncr1}$ mice. (**E**) GO analysis of five distinct macrophage clusters. Dot size represents enriched gene number, and color intensity represents significance. (**F**) Representative flow cytometric plots (left) and cumulative data (right) showing the proportions of KCs and MDMs in total liver cells (**G**) Representative flow cytometric plots (left) and cumulative data (right) showing the relative MFIs of Ly6C in liver MDMs. Data are presented as the mean±SEM and were analyzed by 2-tailed, paired t-test. Differences were evaluated between littermates. Each circle and square on graphs represents an individual mouse; *P*, *P*-value; *, *P*<0.05; **, *P*<0.01, ns, not significant.

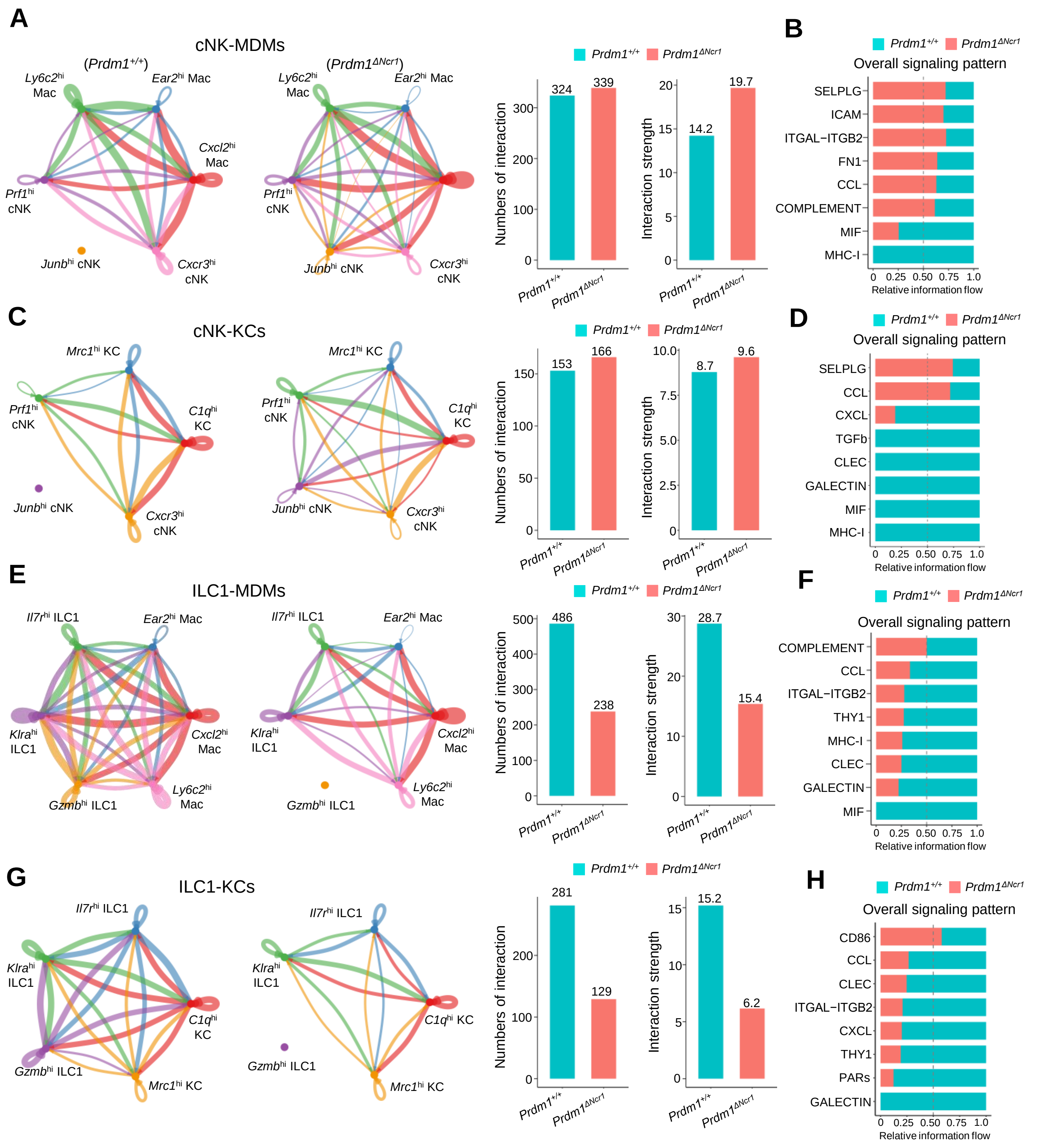


**Supplemental Figure 10. Cell-Cell communication between liver cNK, ILC1s, MDMs, and KCs.**

(**A, C, E** and **G**) Circle plots (left) and summary data (right) illustrating the interaction numbers and strength of significant enriched ligand–receptor pairs in cNK-MDMs (**A**), cNK-KCs (**C**), ILC1s-MDMs (**E**), ILC1-KCs (**G**) interactions between *Prdm1*$^{+/+}$ and *Prdm1*$^{\Delta Ncr1}$ cells. The thickness of the line indicates the number of enrich pairs, and the arrow reflects the direction of the interaction. (B, D, F, and H) Bar graphs showing the information flow in selected active signaling patterns in cNK-MDMs (**B**), cNK-KCs (**D**), ILC1s-MDMs (**F**), ILC1-KCs (**H**) interactions between *Prdm1*$^{+/+}$ and *Prdm1*$^{\Delta Ncr1}$ cells. Relative information flow was calculated as the sum of the communication probability in given signaling patterns.

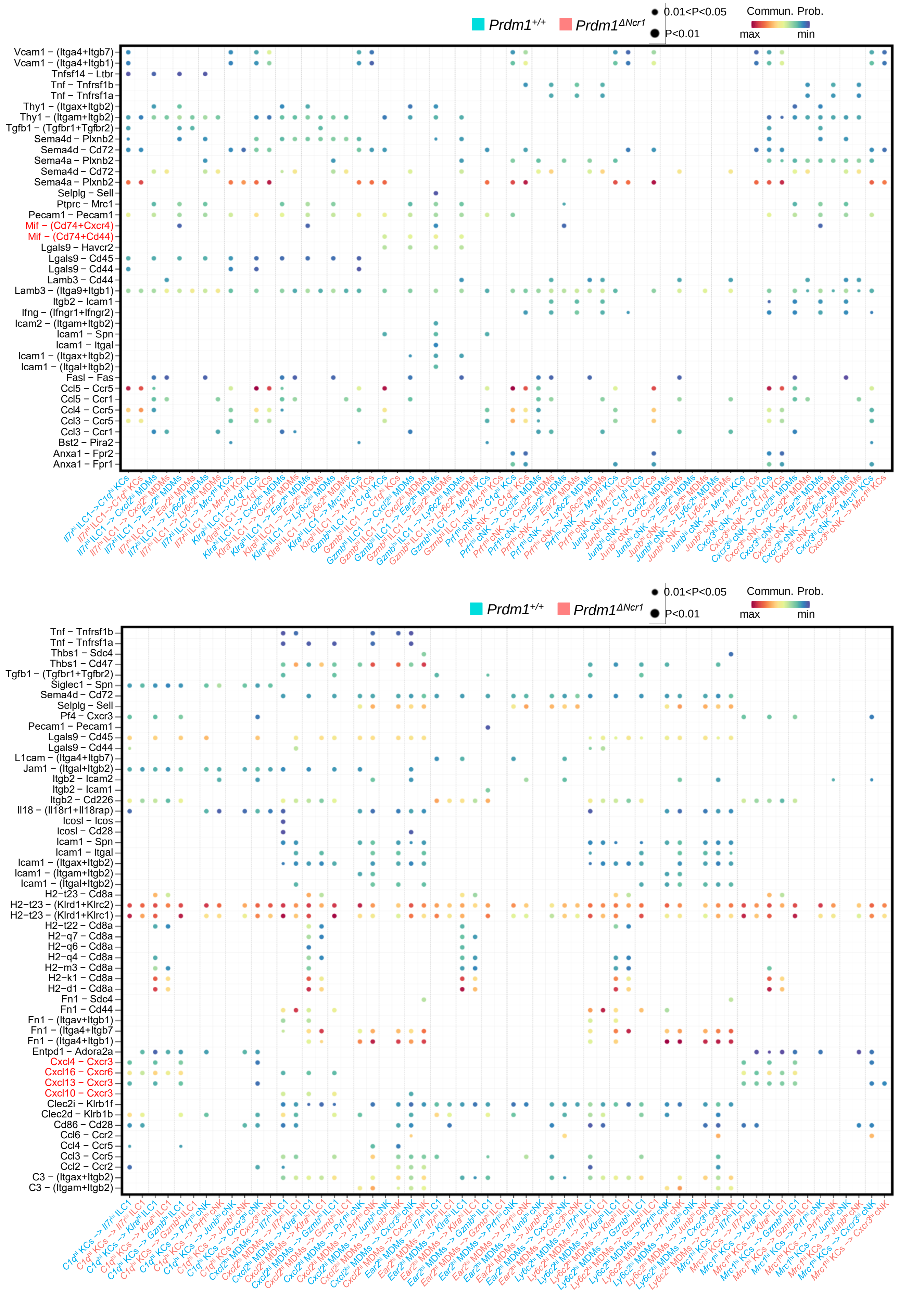

**Supplemental Figure 11. Ligand-receptor interaction between type I ILCs and macrophages.**
Bubble plots showing the significant ligand-receptor pairs between cNK cells, ILC1s, and macrophages within *Prdm1*$^{+/+}$ and *Prdm1*$^{\Delta Ncr1}$ mice. The highlighted Mif-Cd74 and Cxcl-Cxcr signaling was significantly decreased in *Prdm1*$^{\Delta Ncr1}$ mice. Dot size represents the *P*-value, and color intensity represents the communication probabilities. Empty space indicates a communication probability of zero. *P*-value were calculated by the one-sided permutation test.

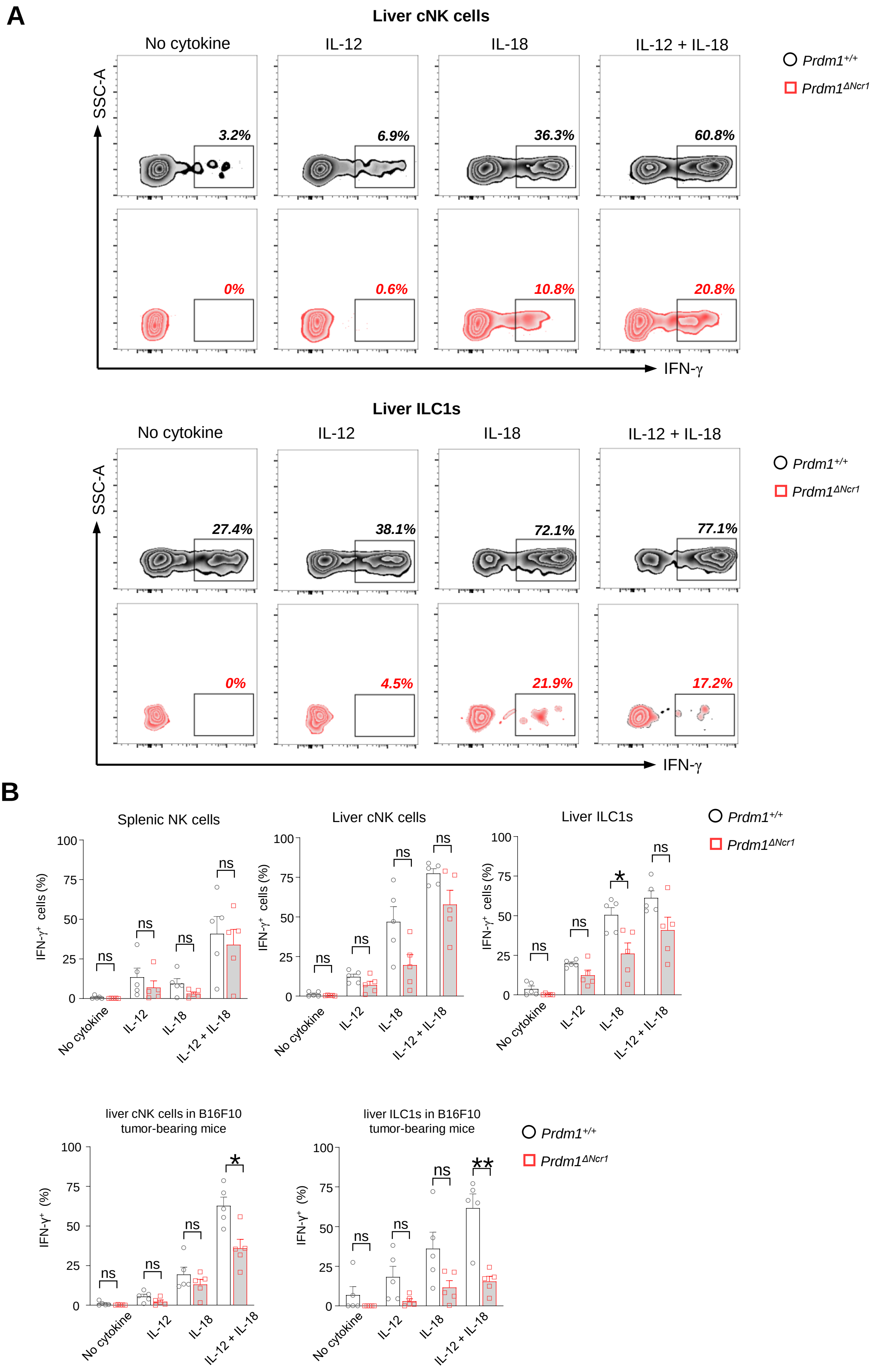


**Supplemental Figure 12:** ***Prdm1*** **maintains the IFN-γ production of liver type 1 ILCs in tumor microenvironment.** (**A**) Representative flow cytometric plot of the frequency of IFN-$\gamma^{+}$ liver cNK cells and ILC1s from *Prdm1*$^{+/+}$ and *Prdm1*$^{\Delta Ncr1}$ tumor-bearing mice at day 14 after inoculation with B16F10 melanoma cells via intrasplenic injection (n=5). (**B**) reanalysis of statistical significance of IFN-γ production in liver cNK cells and ILC1s after IL-12/IL-18 stimulation (Figure 2F and Figure 7E) using unpaired t-test. Data are presented as the mean±SEM and were analyzed by 2-tailed, paired or unpaired t-test. Differences were evaluated between littermates. Each circle and square on graphs represents an individual mouse; *P*, *P*-value; *, *P*<0.05; **, *P*<0.01, ns, not significant.
